# Bioarchaeological sex prediction from central Italy using generalized low rank imputation for cross-validated metric craniodental supervised ensemble machine learning with missing data

**DOI:** 10.1101/2020.11.04.368894

**Authors:** Evan Muzzall

## Abstract

I use a novel supervised ensemble machine learning approach to verify sex estimation of archaeological skeletons from central Italian bioarchaeological contexts with large amounts of missing data present. Eighteen cranial interlandmark distances and five maxillary metric distances were recorded from *n* = 240 estimated males and *n* = 180 estimated females from four locations at Alfedena (600-400 BCE) and two locations at Campovalano (750-200 BCE and 9-11^th^ Century CE). A generalized low rank model (GLRM) was used to impute missing data and 20-fold external stratified cross-validation was used to fit an ensemble of eight machine learning algorithms to six different subsets of the data: 1) the face, 2) vault, 3) cranial base, 4) combined face/vault/base, 5) dentition, and 6) combined cranianiodental. Area under the receiver operator characteristic curve (AUC) was used to evaluate the predictive performance of six constituent algorithms, the discrete algorithmic winner(s), and the SuperLearner weighted ensemble’s classification of males and females from these six bony regions. This approach is useful for predicting male/female sex from central Italy. AUC for the combined craniodental data was the highest (0.9722), followed by the combined cranial data (0.9644), the face (0.9426), vault (0.9116), base (0.9060), and dentition (0.7421). Cross-validated ensemble machine learning of cranial and dental data shows strong potential for estimating sex in the bioarchaeological record and can contribute additional perspectives to help refine our understanding of human sex estimation. Additionally, GLRMs have the potential to handle missing data in ways previously unexplored in the discipline. The main limitation is that the biological sexes of the individuals estimated in this study are not certain, but were estimated macroscopically using common bioarchaeological methods. However, these methods show great promise for estimation of sex in bioarchaeological and forensic contexts and should be investigated on known-sex reference samples for confirmation.

## Introduction

Accurate sex estimation of archaeological skeletal remains is a fundamental step for reconstructing biological and demographic profiles of past humans. After unknown remains are identified, documented, and recovered, the sex and age of deceased individuals and groups are commonly estimated using traditional methods of measuring the pelvis, skull, and teeth (Buikstra & Ubelaker, 1994; Garvin & Ruff, 2012; Krishan et al., 2016). However, because female and male biological maturation rates differ (Slemenda et al., 1994; Wang, 2002), sex misidentification can lead to data recording bias and depreciated interpretability. After sex has been estimated and with the assistance of other biological and archaeological contextual information, the identities and lifeways of the deceased can be reconstructed in bioarchaeological contexts. However, traditional macroscopic sexing methods possess varying degrees of accuracy (Weiss, 1972; Sutter, 2003; Konigsberg, Algee-Hewitt, & Steadman, 2009; Jackes, 2011; Sierp & Henneberg, 2015; Irurita Olivares & Alemán Aguilera, 2016). For example, the dentition, pelvis, and crania can provide different results for estimating sex of the deceased. Tooth crown calcification and eruption and bone epiphyseal fusion are useful until early adulthood when 3rd molars erupt and bony ossification centers fuse in their final, united shapes. However, pelvic and cranial suture methods are used to estimate age in individuals through later stages of adulthood, albeit with wider margins of error.

Craniometric dimensions are frequently used as proxies for genetic relatedness due to their potentially heritable nature and correlations with neutral and adaptive genetic variation and selection (Sjøvold, 1984; Devor, 1987; Roseman, 2004; Roseman & Weaver, 2004; Carson, 2006; Witherspoon et al., 2007; Martínez-Abadías et al., 2009; Strauss & Hubbe, 2010; Herrera, Hanihara, & Godde, 2014). In the absence of genetic information, these methods are used to approximate the genetic and evolutionary relationships of past humans (Buikstra, Frankenberg, & Konigsberg, 1990), thus making accurate sex classification an integral first step in the reconstruction of other biological and demographic parameters. Hence, further examinations of sex correlations with other lines of evidence such as burial location, material culture, occupation, health, trauma prevalence, and biological relatedness will be skewed if sex is first misclassified.

Machine learning has yet to gain a foothold in bioarchaeology despite our discipline’s deep ties to statistics and computational research for investigation of large quantitative datasets. Cunningham’s (1997) pioneering machine learning social anthropological work for rule-based kinship structure detection set a high bar for anthropologists of all subdisciplines to aspire. However, her work remains largely unrecognized even though it exemplifies the types of problem-and-dataset-driven questions faced by bioarchaeologists. This discrepancy persists despite the success of bioarchaeological machine learning applications for estimating sex, age, ancestry, body mass, and stature in forensic anthropology, radiography, and anatomy (Bell & Jantz, 2001; Hefner & Ousley, 2014; Czibula, Ionescu, Miholca, & Mircea, 2016; Ionescu, Teletin, & Voiculescu, 2016; Ionescu, Czibula, & Teletin, 2018; Miholca, Czibula, Mircea, Czibula, 2016; Pink, 2016; Porto et al., 2019; Ortiz, Costa, Silva, Biazevic, & Michel-Crosato, 2020). Even less bioarchaeological research has focused on missing data imputation (Kenyhercz & Passalacqua, 2016).

Therefore, more examples are needed to better test our methodological understandings of skeletal and dental sex estimation. This research is an extension of Muzzall, Kennedy, and Culich (2017) which improved sex prediction accuracy of Howells Worldwide Craniometric Dataset and provided another example of the strong potential for machine learning to assist in sex estimation in bioarchaeological contexts. Here, I use a generalized low rank model to impute large amounts of missing data for a cross-validated supervised ensemble machine learning approach. This framework consists of eight algorithms total and is fit to cranial interlandmark and dental metric distances to predict binary sex from six pelvic and cranially estimated samples at Alfedena (600-400 BCE) and Campovalano (750-200 BCE and 9-11^th^ Centuries CE) in central Italy.

Italy is home to one of the most colossal bioarchaeological contexts on Earth and represents humans’ deep history throughout the region. Its central Mediterranean location and long and complex temporal histories and geological and environmental diversity have been influential in shaping the genetic, morphological, and cultural histories of the region (Scozzari et al., 2001; Coppa, Cucina, Lucci, Mancinelli, & Vargiu, 2007; Muttoni, Scardia, Kent, Swisher, & Manzi, 2009; Fu, Rudan, Pääbo, & Krause, 2012; Ghirotto et al., 2013). Humans here developed some of the richest and most divergent forms of worship, architecture, iconography and writing, and empires that persisted for long periods of time and across the globe via trade, warfare, and colonization. Central Italy was a particular crossroads between Africa and Europe and the Near East and Iberia and was home to many chiefdoms and nation-states that contained both shared and varied forms of settlement patterns, social and burial organization, material cultures, mortuary behaviors, and skeletal-dental morphologies (Muzzall & Coppa, 2019). As a result, Italy’s bioarchaeological record provides a space to experiment with new methodologies for sex estimation.

## Materials and Methods

### Data

The dataset consists of metric cranial and dental data from 240 males and 180 females from central Italy: four locations at the Iron Age necropolis at Alfedena (600-400 BCE), the Iron Age graveyard at Campovalano (750-200 BCE), and the Medieval cemetery at Campovalano (9-11^th^ Centuries CE) (Table 1). Cranial metric data were collected from a total of twelve standard anatomical landmarks: four from the face, four from the cranial vault, and four from the cranial base (Table 2). This produced a total of eighteen cranial interlandmark distances, six from each of the four landmarks from the three cranial regions.

**Table 1.**
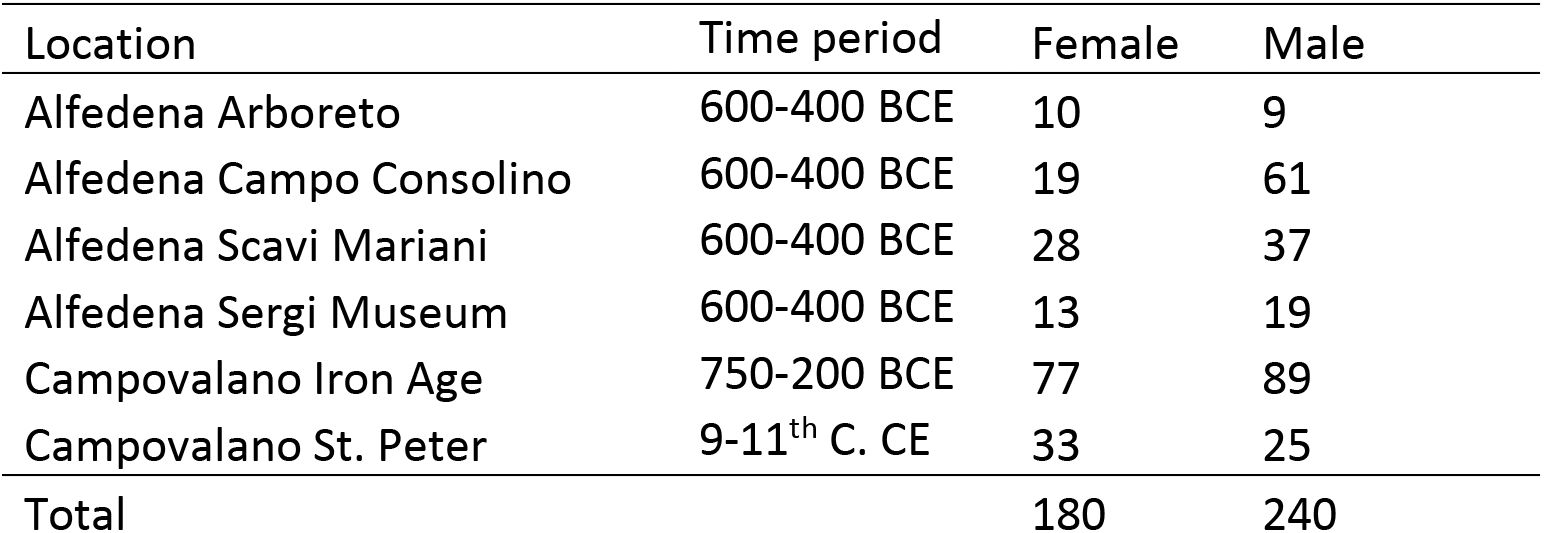
Location, time period, and sex distributions for males and females from Central Italy used in this study.

**Table 2.**
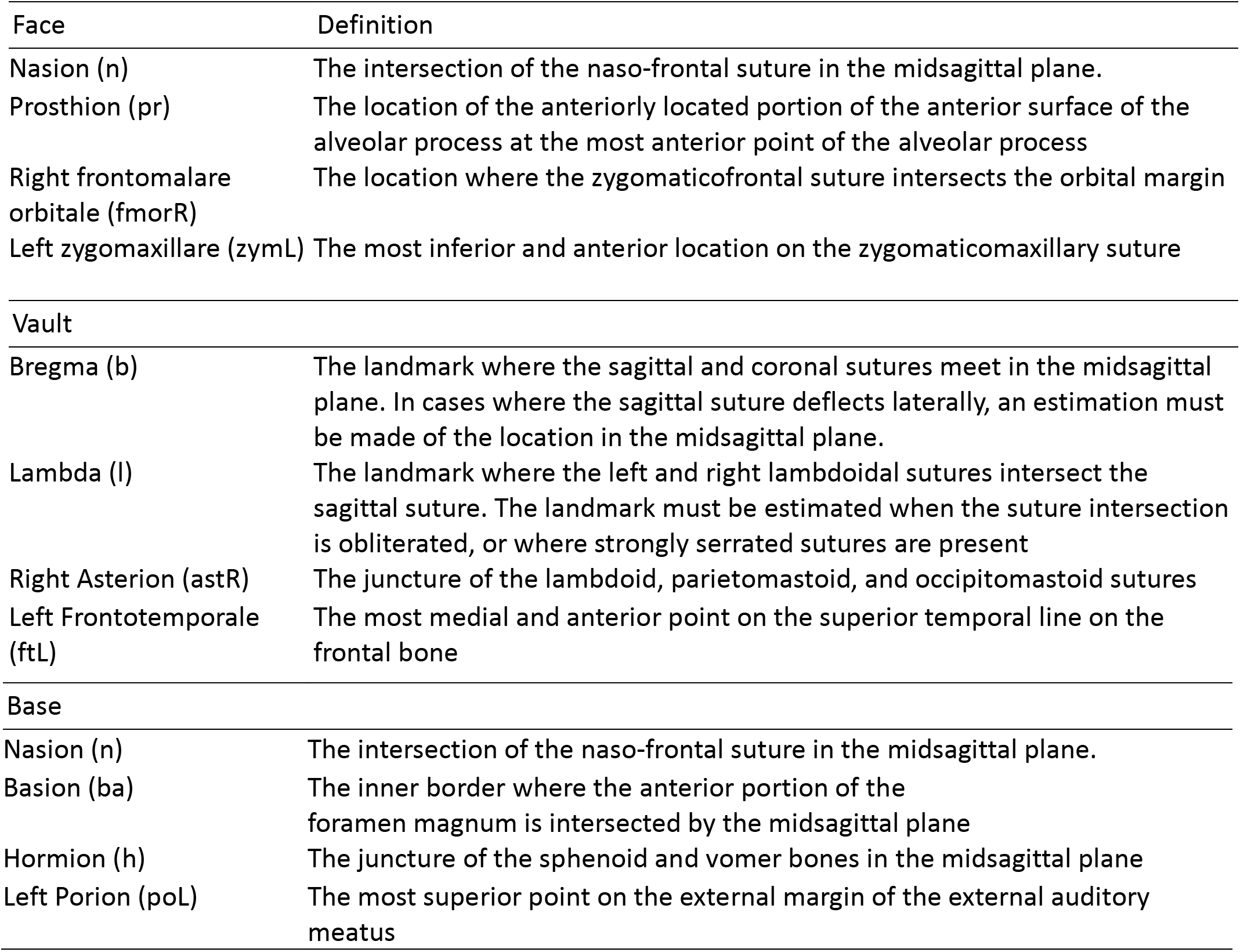
Cranial anatomical landmarks used in this study. The four landmarks from each of the three regions produced eighteen total interlandmark distances – six for each region.

Dental metric data were comprised of maximum mesiodistal dimensions of the right (or left-substituted when right antimere was missing) maxillary canine (XC) and buccolingual breadths of the right mesial (P3) and distal (P4) premolars and first (M1) and second (M2) molars (Hillson et al., 2006). Thus, six different subsets of the data were used: 1) six metrics from the face, 2) six from the vault, 3) six from the base, 4) eighteen from the cranium (the combined face, vault, and base metrics), 5) five from the dentition, and 6) twenty-three from the total combined cranial and dental data. Tukey boxplots are used to illustrate sex differences in these metrics. Sex was originally estimated macroscopically for all samples by Coppa and Macchiarelli (1982) and Bondioli, Corruccini, & Macchiarelli (1986) using methods for the pelvis and crania, except for Campovalano St. Peter which was estimated by EM using discrete cranial traits (Buikstra and Ubelaker, 1994).

### Missing data

Missing data were prevalent from all areas of measurement and percentages of missing values for the face, vault, base, and dentition are shown in Table 3. A generalized low rank model (GLRM) was used to impute the missing values and function as an extension of principal component analysis (PCA) for low rank matrix tabular dataset approximation, by

> “approximating a data set as a product of two low dimensional factors by minimizing an objective function. The objective will consist of a loss function on the approximation error together with regularization of the low dimensional factors. With these extensions of PCA, the resulting low rank representation of the data set still produces a low dimensional embedding of the data set, as in PCA” (Udell, Horn, Zadeh, & Boyd, 2016: 3)

**Table 3.**
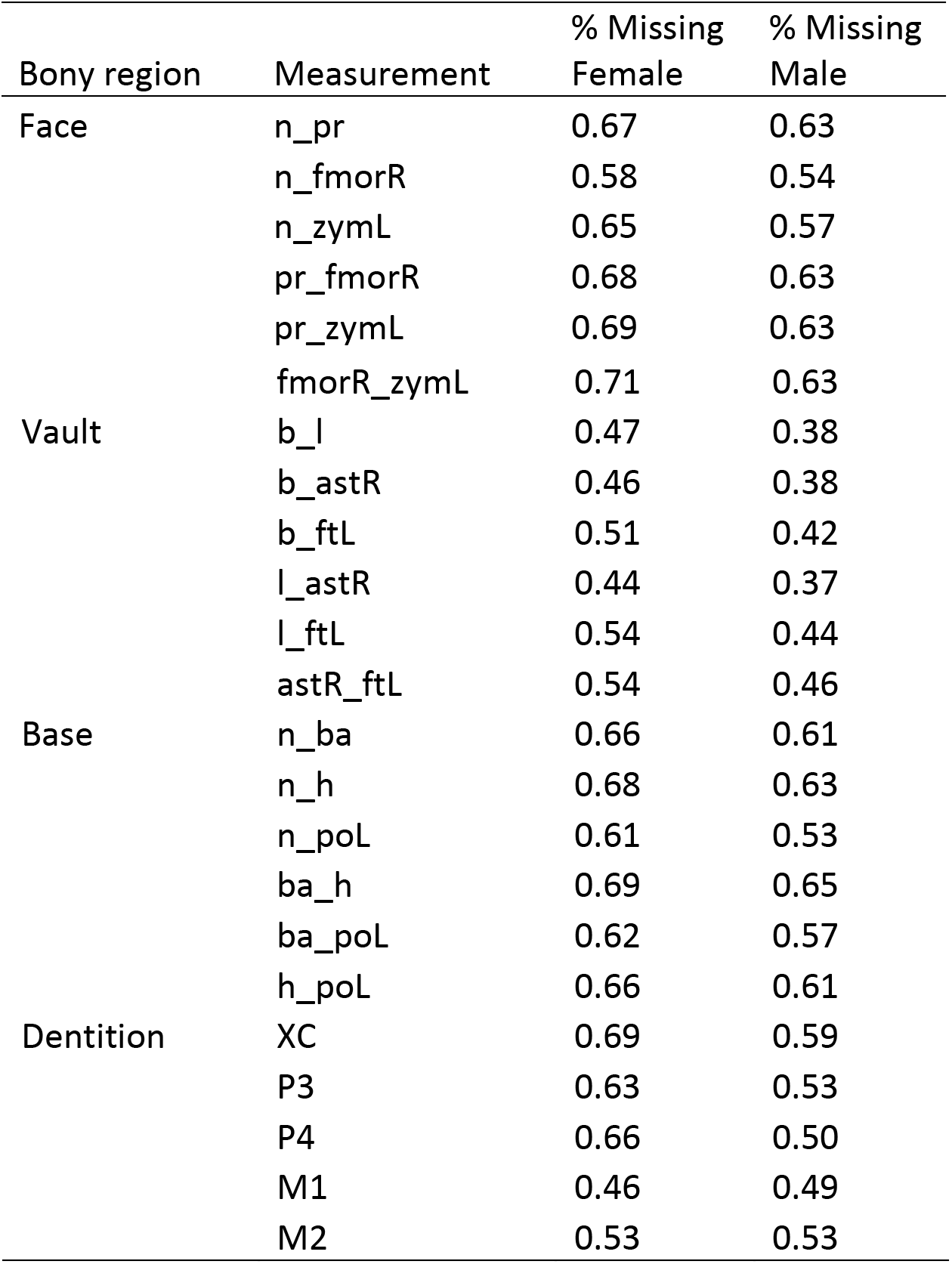
Percentage of missing data for each variable. Missing data were imputed via generalized low rank model.

**Table 4.**
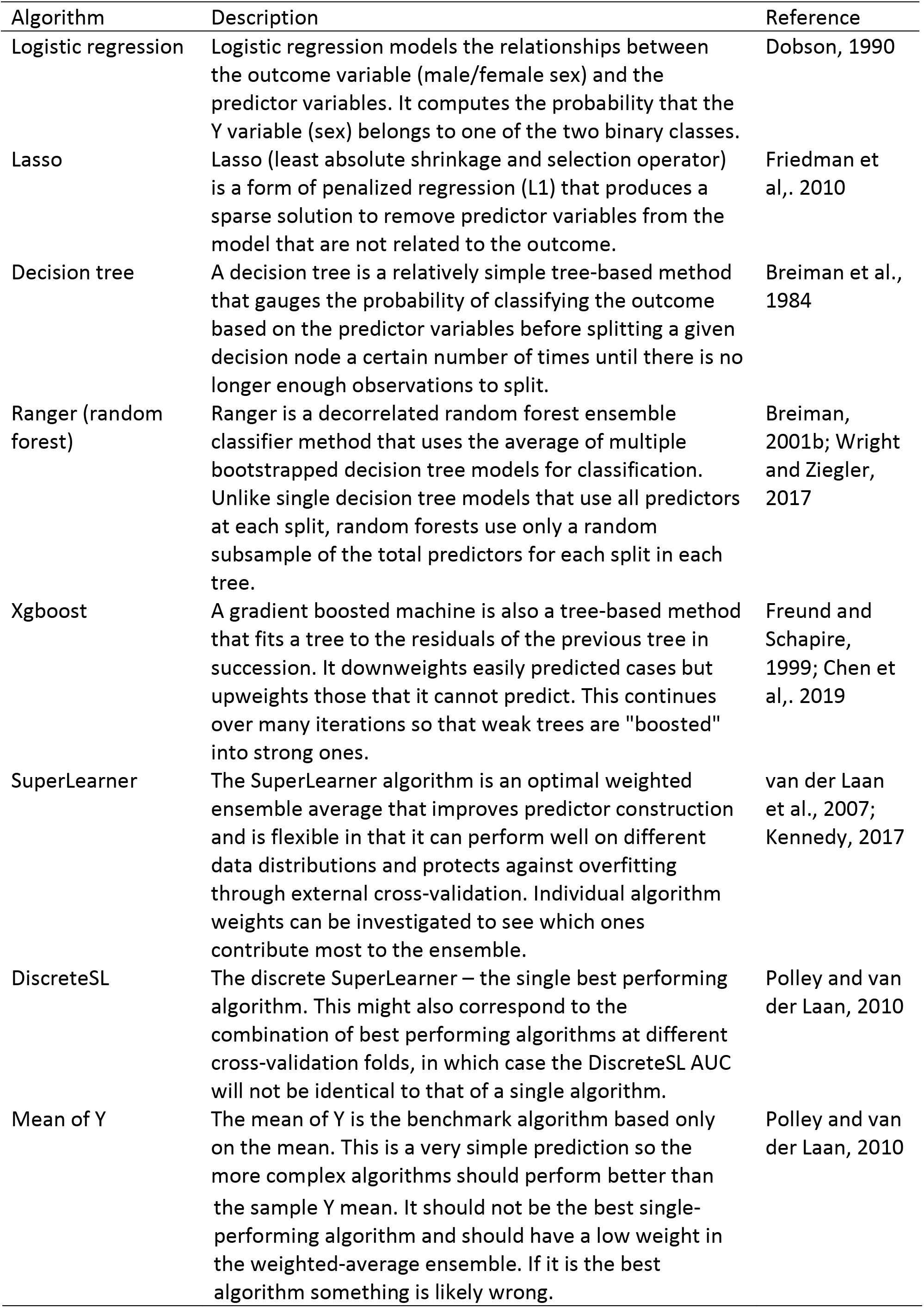
List of eight machine learning algorithms used in this research.

A generalized low rank model is essentially an unsupervised approach for data completion that uses clustering of known data in reduced dimensional space. The advantage of this data-adaptive approach to reconstruct missingness in the skeletal and dental remains instead of column mean, median, or k-nearest neighbor imputation is that it effectively uses clustering of features to impute the missing data, which makes sense given that the missingness of the data arises directly from missingness in the remains. Missingness indicators were also added as columns to the dataset to indicate exactly where missing and imputed data were located. These columns also functioned as predictor variables in the machine learning models to see if the location of missing data was related to sex prediction ability.

### Ensemble Machine learning

Machine learning is defined as “a vast set tools for understanding data” (James, Witten, Hastie, and Tibshirani, 2013:1). It originated as a combination of computer science and statistics, but its greatest strength is its breadth of research application (Breiman, 2001a; Welling, 2015). Early examples stem from the social and cognitive sciences that attempted to predict and imitate human behavior (Turing, 1950; Rosenblatt, 1958; Samuel, 1959). In this research I use a supervised classification machine learning approach because the goal is to predict a categorical outcome (binary male/female sex) using craniodental features as predictor variables.

Ensembles are useful supervised machine learning methods because they optimize predictor accuracy through combinations of a suite of less accurate models (Dietterich, 2000). The SuperLearner approach (van der Laan, Polley, & Hubbard, 2007; Polley & van der Laan, 2010) is an algorithm that uses cross-validation (Efron & Gong, 1982) to estimate the performance of several machine learning models, and/or the same algorithm(s) with different settings. It produces an optimal weighted average of those models (an “ensemble model”), using cross-validated performance. This approach is asymptotically as accurate as the best single prediction algorithm that is tested. I fit the same machine learning ensemble of the eight algorithms (six constituent algorithms, the weighted SuperLearner ensemble, and the single best “discrete” algorithm) to predict binary sex classification (male/female) for each of the six subsets of the data described above as the predictors: face, vault, base, combined cranial, dental, and combined craniodental.

### Evaluating model performance

Stratified cross-validated area under the receiver operator characteristic curve (AUC) was used to evaluate the performance of the individual algorithms and the weighted SuperLearner ensemble (Lantz, 2015; Kennedy, 2017). The receiver operator characteristic curve itself represents the probability that a binary outcome (female or male, in this case) is correctly classified (Hanley and McNeil, 1982) while the AUC provides the degree of separability for the sexes that the model achieves. The receiver operator characteristic curve models the sensitivity (true positive rate) versus specificity (true negative rate) at various thresholds along the receiver operator characteristic curve. Maximization of AUC is ideal, which ranges from 0.5 (equivalent to random guessing) to 1.0 (perfect prediction). AUC is more useful for prediction of imbalanced classes and to prevent overfitting of a single class compared to simple classification accuracy.

Instead of fitting the models separately and looking at the performance (lowest risk), algorithms should be fit simultaneously. Risk is the average loss function used here and measures how far off the prediction was for a given observation and is calculated by nonnegative least squares error; the lower the risk the fewer errors were made by the model. SuperLearner also identifies which single algorithm (or combination of algorithms) is best (the “discrete winner”), in addition to calculating the weighted average of the ensemble itself. Coefficient weights can be viewed to see each algorithm’s contribution to this weighted ensemble average.

Stratified *k*-fold cross-validation is a process that divides the data into equally sized portions and trains a model on *k*-1 portions of the data so that the model can learn the relationship between male/female sex and the various predictors. The one holdout portion is used for testing purposes (but not for fitting the SuperLearner) and this process is repeated *k* times. I chose 20 folds, so each algorithm was trained on 19 portions of the data (95%) and tested on the one holdout (5%), twenty times. This also produces standard errors for the performance of each algorithm that can be compared to the SuperLearner average. Analysis was conducted in R version 3.6.2 and the ck37r, SuperLearner, and ggplot2 packages (Wickham, 2016; Polley, LeDell E, Kennedy, & van der Laan, 2019; Kennedy, 2020).

## Results

Results indicate that ensemble machine learning has strong potential for sex prediction and yielded AUC values greater than 0.90 for the cranial metric data and ~0.74 for the dental metric data, despite large amounts of missing data. Males are larger than females in all dimensions as shown by the Tukey boxplots in Figures 1 and 2 although distributions for the sexes overlap considerably.

**Figure 1.**
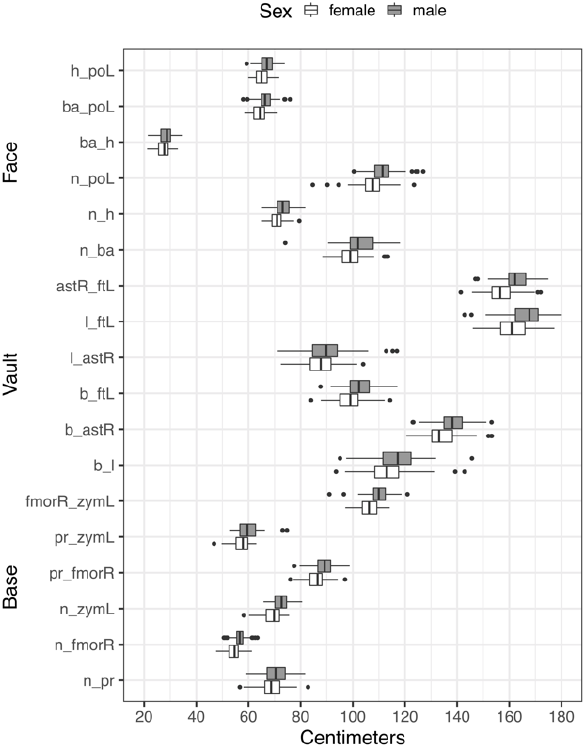
Distributions of raw cranial data for males and females. Anatomical landmark abbreviations are found in Table 2.

**Figure 2.**
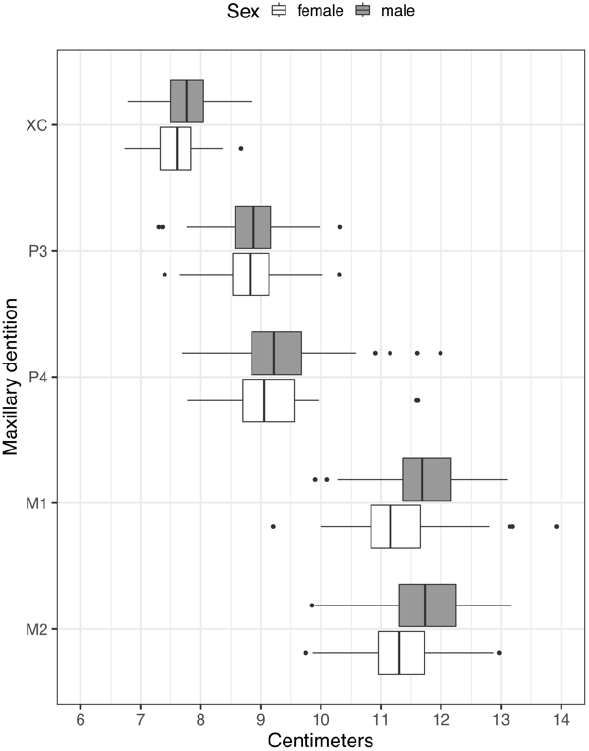
Distributions of raw dental data for males and females. Anatomical landmark abbreviations are found in Table 2.

AUC performance for each algorithm along with their standard errors and confidence intervals are shown in Table 5. The combined craniodental data had the highest AUC with 0.9722, followed by the combined cranial (0.9644), face (0.9426), vault (0.9116), base (0.9060), and dentition (0.7421). Expectedly, the mean of Y is the worst performing algorithm in all cases (AUC = 0.500 for each). The SuperLearner algorithm has the highest AUC for all six bony regions while ranger is a close second for the face, vault, base, cranial, and combined craniodental data. Logistic regression is a close second for the dental data.

**Table 5.**
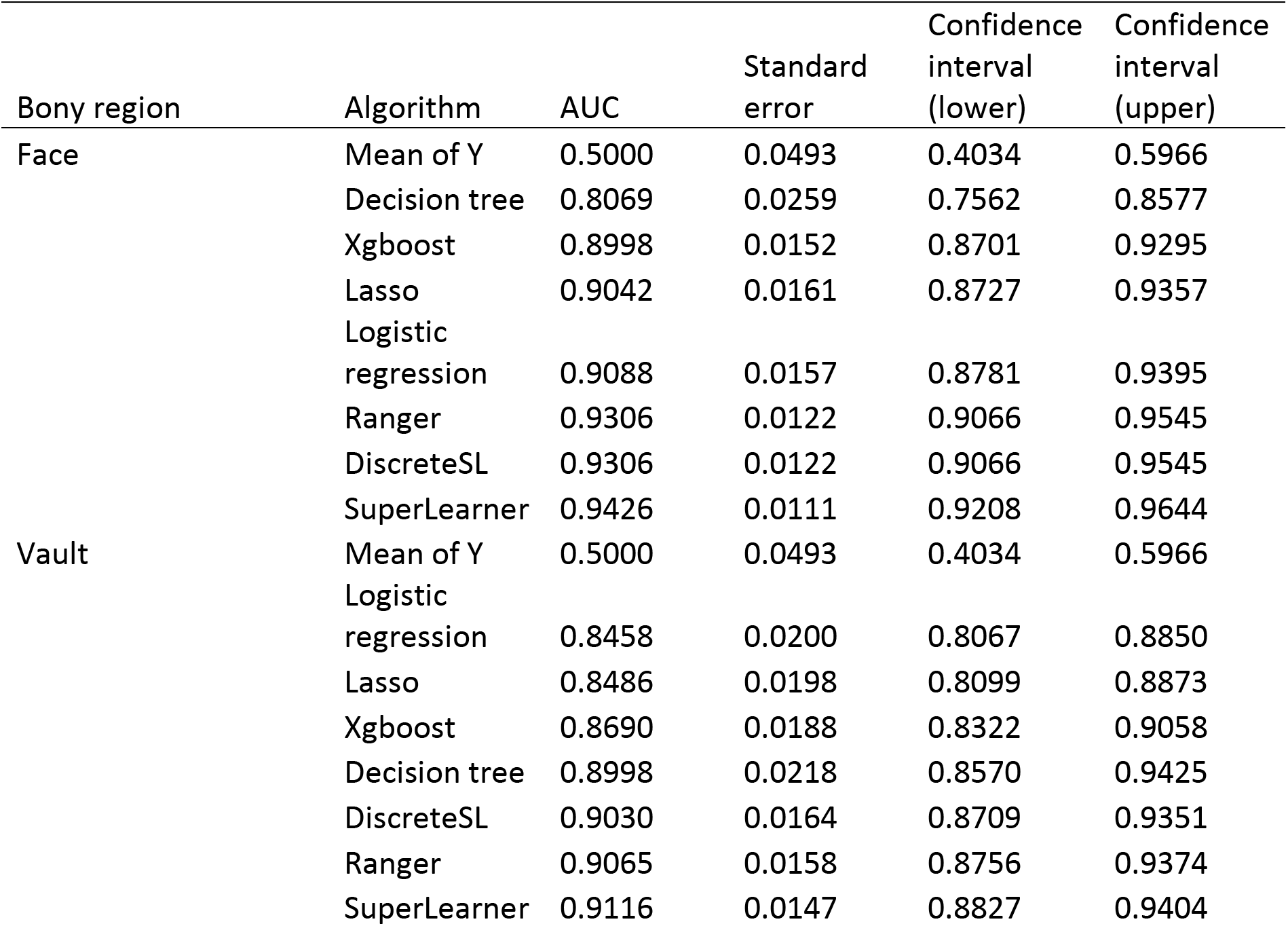
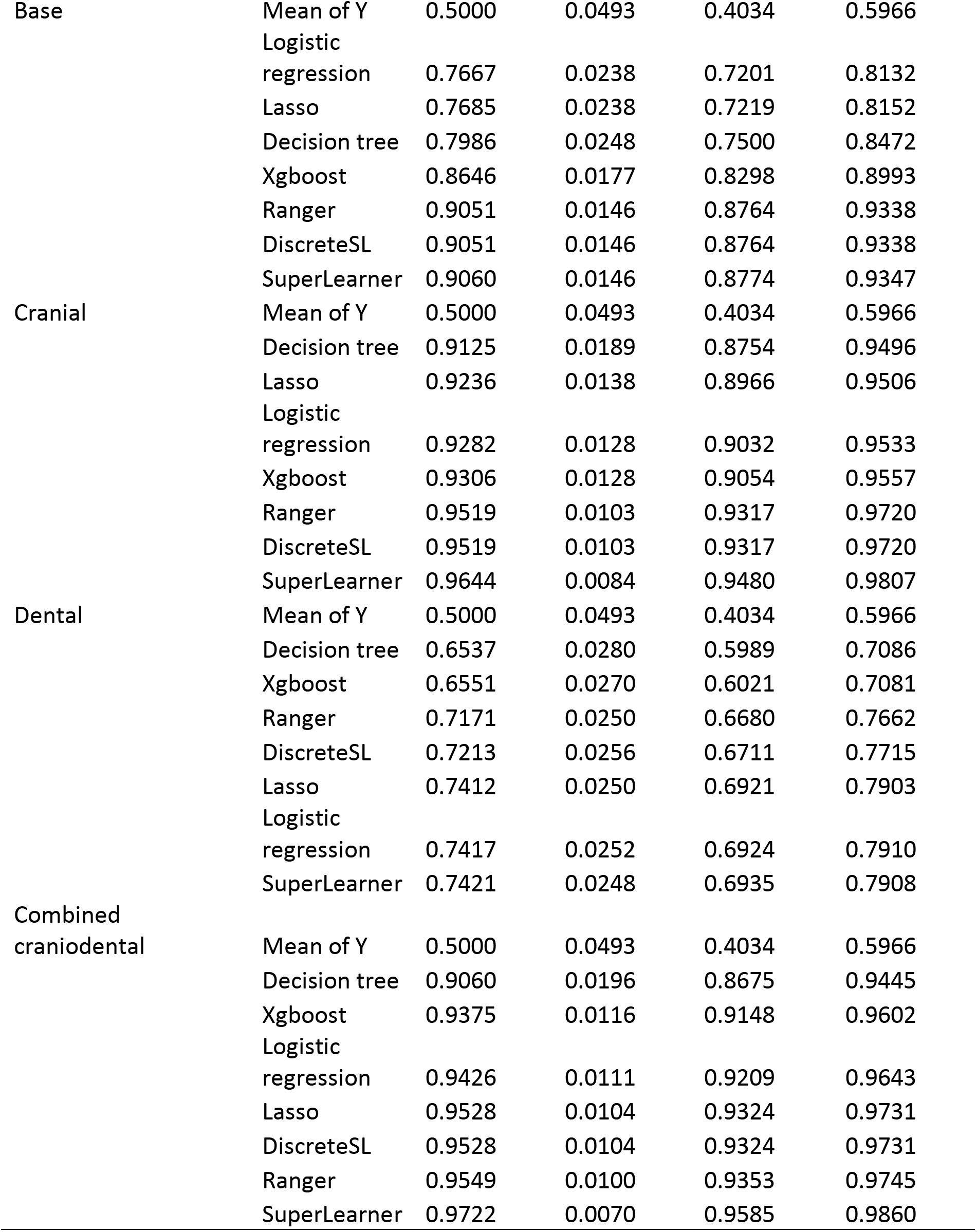
Cross-validated external AUC statistics for the six different measurement regions. 0.5 is the equivalent of random guessing; 1 means perfect prediction.

Also, the single best algorithm (or combination of algorithms) – the DiscreteSL – for the combined craniodental data was the lasso algorithm, with it performing the best for all 20 external cross-validation folds. Ranger was the best-performing algorithm all 20 times for the face, base, and combined cranial data. However, for the vault, ranger was the best performing algorithm 19 times and the decision tree algorithm once. For the dental data, logistic regression was the best performing algorithm 14 times, lasso 4 times, and ranger twice – this algorithmic confusion could be related to the considerably lower AUC for the dentition compared to any of the cranial data.

The SuperLearner weight distributions show which of the individual algorithms contributed most to the ensemble (Table 6). For the combined craniodental data, lasso contributed a coefficient of 0.4522, indicating that it contributed this percentage to the SuperLearner ensemble. This was followed by lesser contributes from the ranger algorithm (0.1734), xgboost (0.1700), logistic regression (0.1319), and decision tree (0.0726). For cranial data, ranger contributed a coefficient of 0.4610, followed by lesser contributions from logistic regression (0.1940), lasso (0.1411), decision tree (0.1267), and xgboost (0.0772). Contributions to the face stem mostly from ranger (0.4634) and logistic logistic regression (0.4193), for the vault from ranger (0.5004) and decision tree (0.3234), and for the base from ranger (0.8878). For the dentition, contributions stem mostly from logistic regression (0.5591) and ranger (0.3582).

**Table 6.**
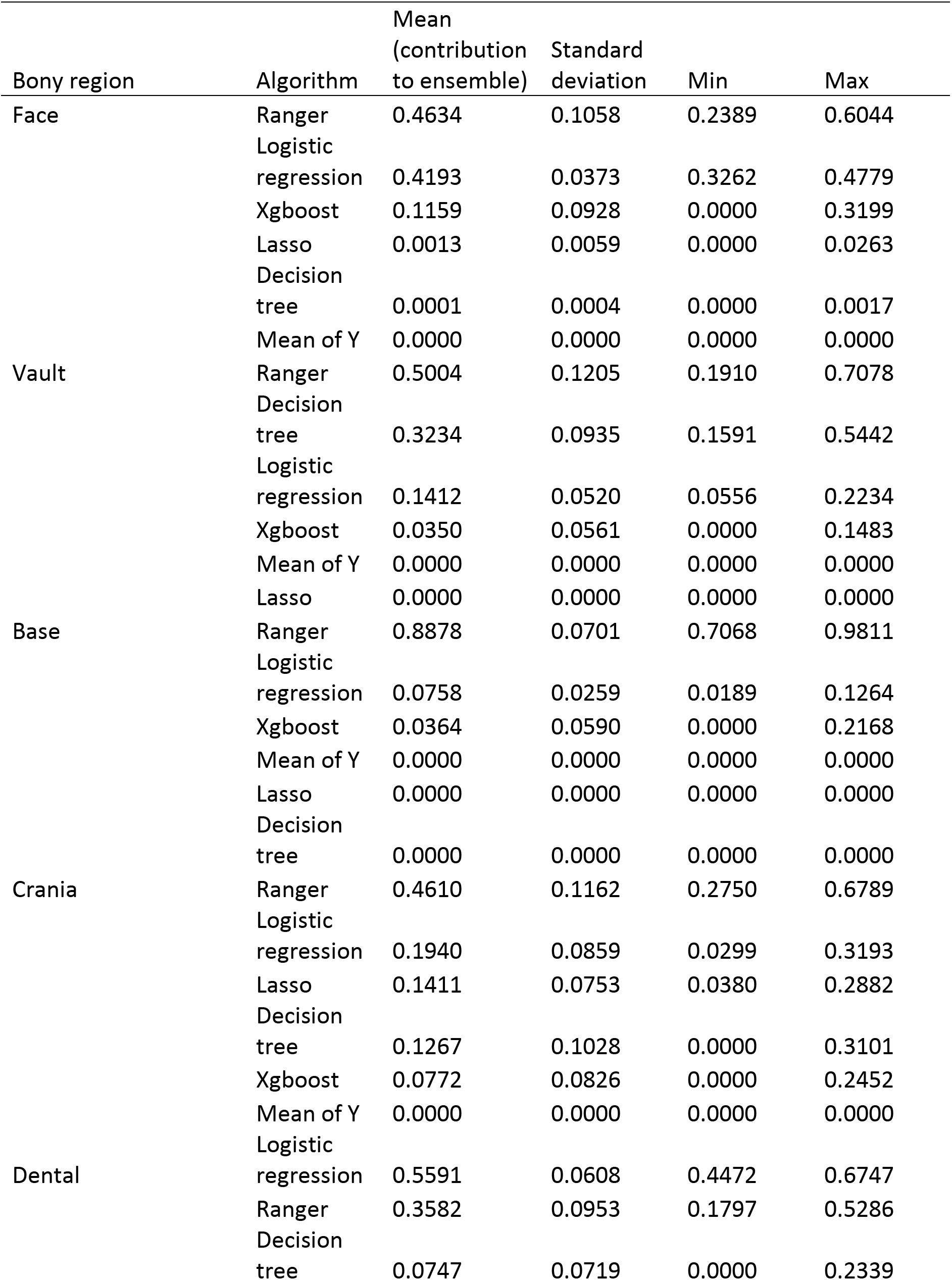
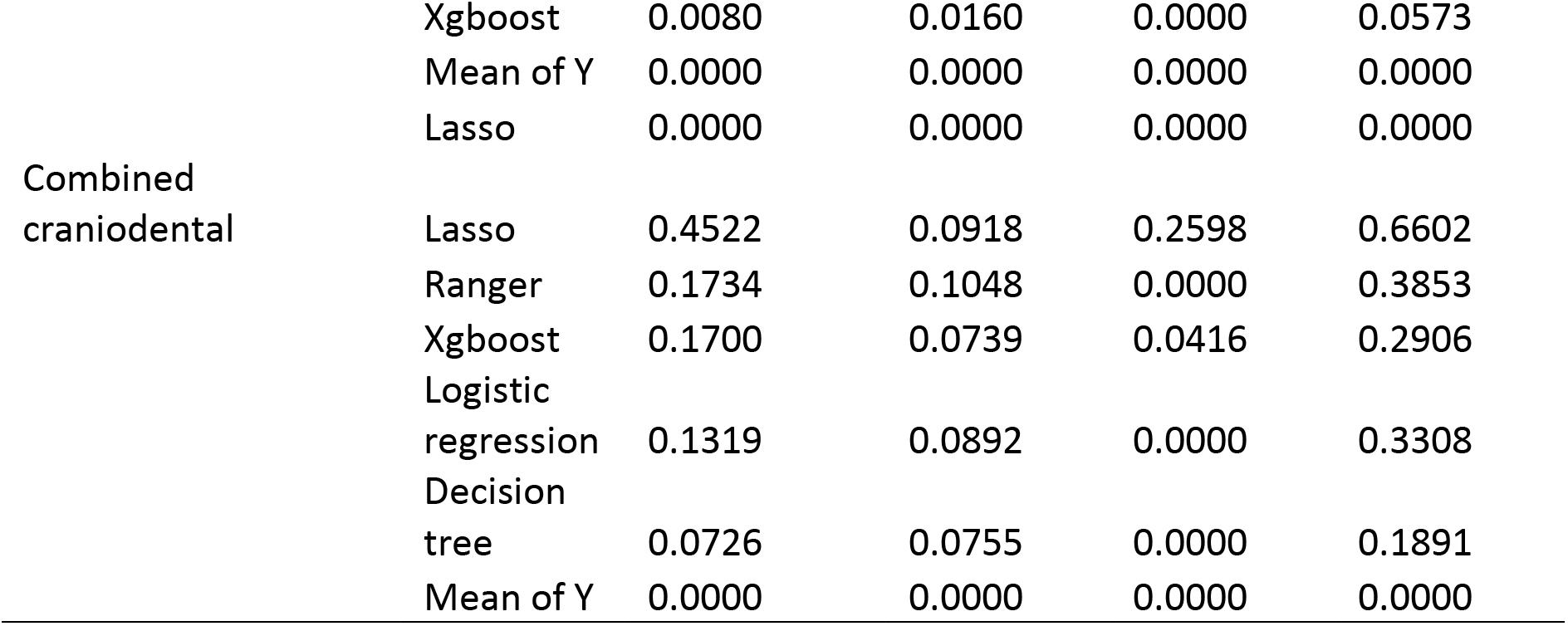
Algorithm weight contributions to the SuperLearner ensembles.

## Discussion

Performance of AUC analysis of the SuperLearner ensemble machine learning framework demonstrates strong potential of this methodology for sex estimation of archaeological remains. An important potential contribution of this research is that it reframes the problem of sex estimation as a predictive one and does not rely on the same assumptions of p-values, traditional hypothesis testing, or causal inference approaches. Additionally, the goal here was not to optimize any algorithms for maximum predictive accuracy, but to instead provide a gentle walkthrough of the process and to stimulate the reader into thinking about how this approach could be applied in their own research contexts. Instead, the focus was on model performance, standard errors, and confidence intervals. This method can also potentially be employed in the field to help resolve disagreements between experts or for indeterminate remains.

Results also support previous research that ensemble machine learning has strong potential for sex estimation in the bioarchaeological record (Muzzall et al., 2017). Although female-ness and male-ness were originally estimated using traits of the pelvis and skull in this study, results support previous research that indicates contrasts between male and female morphological and burial patterns in central Italy during the Iron Age (Coppa & Macchiarelli, 1982; Bondioli et al., 1986; Rubini, 1996; Muzzall & Coppa, 2019). Of particular interest was the general size differences between males and females despite their overlapping distributions. If the modeling process was strongly influenced by size however, it would be reasonable to expect that the dentition would have higher AUC values similar to that of the cranial data. Whether or not the antimeric substitution of left teeth for right teeth in the absence of a rightside tooth and/or the sheer amount of missingness influenced the much lower dental AUC is unknown. More cranial-dental comparisons are necessary to evaluate the reliability of the dentition in this framework. Among the three different cranial regions, the face had the highest AUC values, followed by the base and vault. This could provide further support of the utility of the face for population reconstruction despite its greater environmental plasticity compared to the base and vault due to sensory functions of sight, smell, and taste (Taubadel, 2009).

The ensembles themselves can be strengthened by including more algorithms and customizing them with varying hyperparameters (pre-training settings) to find the most accurate and best performing tunings (Bergstra & Bengio, 2012). Other considerations can be more thoroughly incorporated as well, such as different confusion matrix derivations to evaluate performance such as precision and recall to help highlight class imbalance problems, balanced estimator constructions, false discovery rate, and F1 score. Negative log-likelihood could also be used as the optimizer instead of nonnegative least squares. Other algorithms and methods might be more appropriate – only 6 algorithms with default settings were incorporated in this project but many others can be included in the ensemble (e.g., Bayesian additive regression trees, Chipman; George, & McCulloch, 2010). Features can be screened to identify more interpretable models and custom algorithms can be included to the researcher’s exact specifications (see Kennedy, 2017 for the walkthrough in R). Moreover, deep learning – a subdiscipline of machine learning that utilizes multi-layered artificial neural networks for modeling, predicting, and understanding data – might be even more (Chollet & Allaire, 2017). When dataset sizes and the number of algorithms exceed personal compute potential, the software packages for analyses mentioned in this research have instructions to be run in parallel across multiple cores on a single computer or across multiple machines in cluster or remote settings. Perhaps of great interest to the bioarchaeologist, variable importance information can be extracted from the tree-based algorithms to see which cranial and dental dimensions have the highest weights for sex classification.

It is critical to note that due to the antiquity of the samples included in this research, the sexes of the individuals utilized were estimated macroscopically using features of the pelvis and skull and that the sexes were not actually certain thus making this study a sort of “estimation of estimations”. Known-sex references samples should be a prerequisite for confirmation of methods presented here, and larger sample sizes might be important as well. This study is merely a demonstration of the methods and advertisement of the potential forgeneralized low rank imputation and ensemble machine learning processes in bioarchaeological and forensic contexts. Cadaver samples and skeletal collections would be particularly useful for testing procedures outlined here.

Ensemble machine learning techniques should be considered as part of the bioarchaeologist’s toolkit as an additional method for comparison to macroscopic interrogations of the skeleton and dentition that we rely upon for reconstruction of the biological profiles of past humans. These techniques can potentially assist not only in bioarchaeological reconstructions, but also in forensic applications for identification of missing persons and perhaps even to material, faunal, and floral assemblages as well as mortuary studies and settlement organization. Furthermore, GLRMs warrant further exploration and should be considered by bioarchaeologists as a potentially strong data preprocessing tool when faced with missing data and analytical techniques that require full datasets for computation. Social scientists in general would benefit from updating their instrumentation with crossvalidated ensemble machine learning techniques when research requires some variable(s) to be predicted.

## Acknowledgements

I gratefully thank the UC Berkeley D-Lab, Alfredo Coppa, Chris J. Kennedy, and the Soprintendenza Archeologia d’Abruzzo, and the staffs from the Museo Antropologia de “Giuseppe Sergi” – Sapienza, Museo Paludi di Celano, Museo Archeologico Nazionale d’Abruzzo di Chieti, and Museo di Archeologico Nazionale di Campli. Aniket Kesari and Patrick Muzzall provided comments that improved the quality of this manuscript.

## Notes

### Competing Interest Statement

The authors have declared no competing interest.

### Summary of Updates

Revision 3

